# poRe: an R package for the visualization and analysis of nanopore sequencing data

**DOI:** 10.1101/007567

**Authors:** Mick Watson, Marian Thomson, Judith Risse, Richard Talbot, Javier Santoyo-Lopez, Karim Gharbi, Mark Blaxter

**Affiliations:** Edinburgh Genomics, The Roslin Institute and R(D)SVS, University of Edinburgh, Easter Bush, EH25 9RG; Edinburgh Genomics, Ashworth Laboratories, University of Edinburgh, W Mains Rd, Edinburgh, EH9 3JT

## Abstract

**Motivation:** The Oxford Nanopore MinION device represents a unique sequencing technology. As a mobile sequencing device powered by the USB port of a laptop, the MinION has huge potential applications. To enable these applications, the bioinformatics community will need to design and build a suite of tools specifically for MinION data.

**Results:** Here we present poRe, a package for the statistical software R that enables users to manipulate, organize, summarise and visualize MinION nanopore sequencing data. As a packge for R, poRe has been tested on both Windows and Linux. Crucially, the Windows version allows users to analyse MinION data on the Windows laptop attached to the device

**Availability:** Pre-built R packages for Windows and Linux are available under a BSD license at http://sourceforge.net/projects/rpore/

## 1 INTRODUCTION

Relative to first- and second- generation sequencing technologies, single molecule sequencing (SMS) is a new science, with only Helicos (Bowers, et al., 2009), Pacific Biosciences (Eid, et al., 2009) and Oxford Nanopore (ONT) being widely available. Even within the field of SMS, ONT’s nanopore sequencing technology represents a new paradigm; whilst both Helicos’ and Pacific Biosciences’ sequencing technologies measure incorporation events into a second strand, ONT’s MinION and GridION systems measure a single molecule of DNA as it passes through a protein nanopore. In addition, the MinION is the world’s first mobile DNA sequencer; it is powered by a laptop’s USB port and measures approximately 6” in length. Recently, ONT opened up the MinION access programme, enabling researchers to use the device for the first time.

The ultra-low-cost, mobile nature of the MinION device opens up a huge number of applications. However, users of the device are faced with a number of informatics challenges. Users of the MinION must buy a high-specification Windows laptop, and thus there is a need for Windows-based software to handle the data. The MinION outputs binary files in the HDF5 format (http://www.hdfgroup.org/HDF5/). These contain raw data from the sequencer, which are then processed by a cloud-based base-caller called “metrichor”. The subsequent called sequence files are also in HDF5 format (with the extension .fast5). It is not uncommon for users to be presented with 30-50,000 HDF5 files (.fast5), with no software with which to access the data. Furthermore, data from all runs are stored in a single directory, with no sub-directories, and users find themselves needing to manipulate thousands of files manually, which takes time and is error-prone.

We have developed poRe, a package for the statistical package R, which enables users to manipulate MinION fast5 files into run folders, extract fastq, gather statistics on each run and plot a number of key graphs such as read-length histograms and yield-over-time. Crucially, as a package for R, poRe runs on both Windows and Linux. The Windows version enables users to run poRe on the MinION laptop itself, rather than copying the data to a Linux server to process with Perl or Python. This key feature brings users closer to true mobile DNA sequencing.

## 2 METHODS

### 2.1 Data format

The fast5 HDF5 files contain a number of hierarchical groups, datasets and attributes. An example structure is:

~~~
/Analyses
        /Basecall_2D_000
                 /BaseCalled_2D
                 /BaseCalled_template
                 /BaseCalled_complement
                 /Configuration
                 /HairpinAlign
                 /Summary
        /EventDetection_000
                 /Reads
/Sequences
/UniqueGlobalKey
~~~

The raw events reported by each nanopore, as well as fastq data called by the metrichor base-caller, are embedded within the /Analyses/Basecall_2D_000 group, whereas meta-data about the run itself is embedded within the /UniqueGlobalKey group.

### 2.2 Organisation

The first task users face is to organize a single MinION folder, which may contain reads from many different runs. We provide the function *copy.runs()* to help with this. The function reads all fast5 files within a user-defined directory, and extracts both the unique run identifier (“run_id”) and the name and version of the base-caller. Each read is then copied to a user-defined destination folder, under sub-folders defined by the run_id and the name and version of the analysis. The latter is key, as each raw read may be base-called many times by different versions of the metrichor base-caller

### 2.3 Fastq extraction

Once the data are organized, users may wish to extract fastq data. This can be done using the *extract.run.fastq()* function. For each fast5 read in a given directory, this function will extract the template, complement and 2D fastq data where they exist, and write these to individual fastq files.

### 2.4 Run statistics

Embedded within each fast5 file are a number of key statistics about the reads, such as read length, the number of events, the time of the analysis, mean quality scores, device ID, asic ID, asic temperature and channel number. These can be extracted for all reads in a run by the function *read.fast5.info()*. This returns a data frame with rows as reads and 24 columns of meta-data for each read. Once this data frame has been extracted, the function *run.summary.stats()* can be used to extract key summary statistics, such as max, min and mean read lengths.

### 2.5 Data exploration

We provide a number of functions that allow users to explore the data visually. A feature of all of the visualization functions is that, not only do they plot to the current device, but where appropriate they return data to the user in the form of a data frame, so that the user can then create their own plots in, for example, using ggplot.

Histograms of read length can be created using the *plot.length.histogram()* function. This plots histograms for the template, complement and 2D read lengths, and an example can be seen in Figure 1.

**Figure 1.**
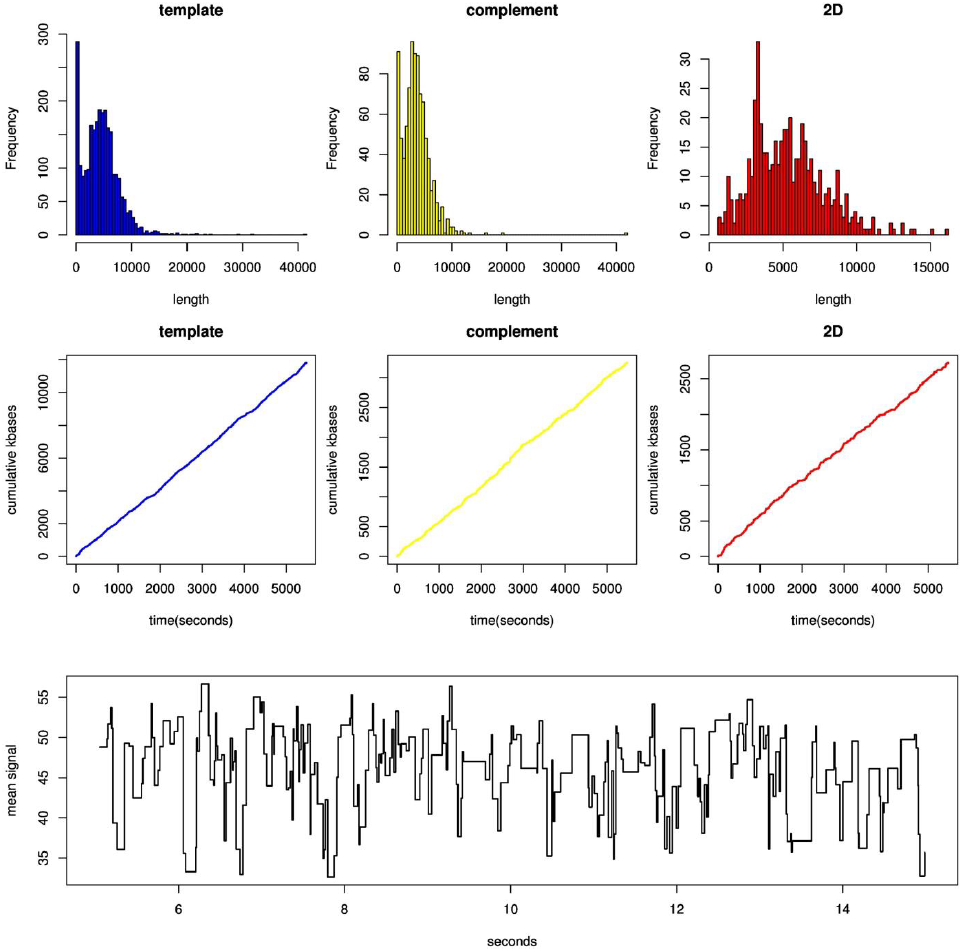
Example plots from poRe showing read length histograms (top), cumulative yield over time (middle) and events from nanopore sensing of a DNA molecule (bottom)

The *plot.cumulative.yield()* function can be used to plot cumulative yield of the run over time, and sums up the template, complement and 2D read lengths over time in seconds since the analysis began. An example of this can also be seen in Figure 1.

Finally, the MinION device consists of a number of channels, each of which should contain a single nanopore. Users can count and plot the number of reads per channel for a run, using *plot.channel.reads()*, and sum and plot the yield per channel, using *plot.channel.yield().* Both of these can be potentially used to diagnose problems in particular areas of the flowcell

### 2.6 Extracting and plotting events

The raw data from the MinION is information about the electronic signal measured as each single molecule of DNA passes through the protein nanopore. It is this data that is converted to sequence data by the metrichor agent. However, the raw events data is also available and can be extracted using the function *get.events()*. This will extract the thousands of events for both the template and complement for a particular read.

The events data may then be visualized using the *plot.squiggle()* function, an example of which can be seen in Figure 1.

## 3 DISCUSSION

We have written poRe, an R package that enables users to more easily manipulate, summarise and visualize MinION nanopore sequencing data. As a package for R, poRe is available for both Windows and Linux, and crucially the Windows version will allow data analysis on the mandatory Windows laptop on which the MinION depends. In addition, R is now a popular statistical package amongst biologists, who may feel comfortable using poRe through the R user-interface.

poRe is one of the first bioinformatics packages to offer this necessary functionality. `poretools` (Loman and Quinlan, 2014) a toolkit written in Python, offers similar functionality, although each software has a different set of (overlapping) functions. We do not believe that `poretools` has been tested on the Windows operating-system. The cross-platform nature of poRe, in addition to poRe’s ability to organize folders of fast5 files, makes poRe an important tool for users of the MinION.

## ACKNOWLEDGEMENTS

We would like to thank Oxford Nanopore for granting Edinburgh Genomics access to the MinION Access Programme (MAP)

### Funding

Edinburgh Genomics is partly supported through core grants from NERC (R8/H10/56), MRC (MR/K001744/1) and BBSRC (BB/J004243/1).

